# Mutation accumulation in selfing populations under fluctuating selection

**DOI:** 10.1101/230326

**Authors:** Eddie K.H. Ho, Aneil F. Agrawal

## Abstract

Selfing species are prone to extinction, possibly because highly selfing populations can suffer from a continuous accumulation of deleterious mutations, a process analogous to Muller’s ratchet in asexual populations. However, current theory provides little insight into which types of genes are most likely to accumulate deleterious alleles and what environmental circumstances may accelerate genomic degradation. Here we investigate temporal changes in the environment that cause fluctuations in the strength of purifying selection. We simulate selfing populations with genomes containing a mixture of loci experiencing constant selection and loci experiencing selection that fluctuates in strength (but not direction). Even when both types of loci experience the same average strength of selection, loci under fluctuating selection contribute disproportionately more to deleterious mutation accumulation. Moreover, the presence of loci experiencing fluctuating selection in the genome increases the deleterious fixation rate at loci under constant selection; under most realistic scenarios this effect of linked selection can be attributed to a reduction in *N_e_*. Fluctuating selection is particularly injurious when selective environments are strongly autocorrelated over time and when selection is concentrated into rare bouts of strong selection. These results imply that loci under fluctuating selection are likely important drivers of extinction in selfing species.

## Introduction

Self-fertilization has evolved many times independently in plants and animals (Goodwillie et al. 2005, Jarne and Auld 2006). Despite frequent transitions from outcrossing to self-fertilization, highly selfing populations are relatively rare in nature and relegated at the tips of phylogenies (Stebbins 1957, Igic and Kohn 2006). Stebbins (1957) proposed that highly selfing species suffer from low rates of adaptation and are more prone to extinction; classically called the ‘dead end’ hypothesis. Population genetics theory provides insights into why selfing is a dead end. Selfing reduces the effective population size, *N_e_*, which diminishes the efficacy of selection. Under neutrality, *N_e_* is reduced two-fold in obligately selfing populations because gametes are not sampled independently (Pollak 1987). In addition, but perhaps more importantly, high homozygosity in selfing populations reduces the effectiveness of recombination and can further reduce *N_e_* due to linked selection effects (Nordborg 1997, Roze 2016). Explicit contrasts of selfing and outcrossing species confirm lower rates of adaptation and higher rates of deleterious mutation accumulation in both theory (Glemin and Ronfort 2013; Kamran-Disfani and Agrawal 2014) and reality (Slotte et al. 2010, Arunkumar et al. 2014, Burgarella et al. 2015). Further, phylogenetic analyses infer higher extinction rates in self-compatible *Solanacaea* (Goldberg et al. 2009) relative to their self-compatible counterparts. The disadvantage resulting from very strong linked selection can maintain low levels of outcrossing in species considered strong selfers (Kamran-Disfani and Agrawal 2014).

While both reduced rates of adaptation and higher rates of deleterious mutation accumulation may contribute to extinction, our focus here is on better understanding the latter. Though existing theory makes it clear that deleterious mutations accumulate more readily in highly selfing populations, this literature does not aim to provide insight into the types of genes most likely to degrade or the types of environmental circumstances that might hasten genomic degradation. Here we investigate how loci that experience fluctuations in intensity (but not direction) affect deleterious mutation accumulation in highly selfing populations. As described below, fluctuations in selection intensity affect fixation probabilities in both outcrossing and selfing species but we focus on the latter because deleterious mutation accumulation is, in general, expected to be much more prevalent in selfers.

There is abundant evidence that selection is environmentally sensitive in natural and in laboratory experiments (Leimu and Fischer 2008, Bergland et al. 2014, Latta et al. 2015, Rutter et al. 2017). Conceptually, selection can vary in direction and magnitude between environments (Martin and Lenormand 2006), but empirically, selection varies mostly in magnitude. For example, ‘stressful’ environments may alleviate or intensify the strength of selection, but rarely reverses the direction of selection (Kishony and Leibler 2003, Agrawal and Whitlock 2010, Kraemer et al. 2015). Hillenmeyer et al. (2008) found that 97% of all viable singe gene knockouts in yeast reduce fitness in at least one environment while having little effect in other environments after performing over 1000 chemical assays. This suggests that selection against mutations can fluctuate in magnitude from neutrality to deleterious between environments (i.e., conditionally neutral mutations may be common). Other types of studies also indicate that temporal changes in the environment create fluctuations in the strength of selection (Grant and Grant 2002, Bell 2010, Siepielski et al. 2017). Despite this evidence, most mutation accumulation models assume selection is constant. This likely underestimates deleterious fixation rates if selection fluctuates temporally (Wardlaw and Agrawal 2012, Cvijovic et al. 2015).

Though classic single locus models for the fixation probability assume selection is constant (Haldane 1927, Kimura 1962), fixation probabilities are known to be highly sensitive to selection changing over time (Kimura and Ohta 1970, Gillespie 1978, Peischl and Kirkpatrick 2012). Often neglected are changes in selection that are stochastic and temporally autocorrelated, especially for mutations that are on average deleterious. Environments can fluctuate over many different time scales under various degrees of temporal autocorrelation (Halley 1996, Vasseur and Yodviz 2004, Furguson et al. 2016) and evolutionary processes can be strongly dependent on both the temporal autocorrelation in selection and the average strength of selection (Charlesworth 1993, Pieschl and Kirkpatrick 2012, Wardlaw and Agrawal 2012). In general, if changes in selection occur rarely (very high temporal autocorrelation), then the fixation probability for the mutation is largely dependent on the selective environment in which it arose and can be estimated from models assuming constant selection. If fluctuations are very frequent (very low temporal autocorrelation), fixation probabilities can also be approximated by constant selection models but using a time-averaged strength in selection. However, in the intermediate range where selection and environmental change operate on similar time scales, classic equations become inaccurate. Within this range, Cvijovic et al. (2015) have shown in single locus models that selection fluctuating in direction increases the probability of fixation for all mutations.

We examine selection that is always negative (or zero) but fluctuates in intensity (but not direction) in a system with strong linked selection due to selfing. Interference between deleterious mutations leads to mutation accumulation in asexual and highly selfing populations (Muller 1932, Haigh 1978, Charlesworth and Charlesworth 1993, 1997, Kamran-Disfani and Agrawal 2014). However, little is known about selective interference when some loci experience fluctuations in selection intensity. In a related study, Wardlaw and Agrawal (2012) found that temporal autocorrelation in selection intensity is an important determinant of Muller’s ratchet in asexuals because the ratchet can turn quickly during periods of relaxed selection. The current work extends their findings to selfing populations with some recombination, explores more realistic environmental scenarios, separately quantifies fixation rates for loci experiencing fluctuating versus constant selection within the same genome, examines the extent to which the linked selection effects from fluctuations can be accounted for through *N_e_*, and considers selective interference involving beneficial mutations.

Our goal here is to investigate mutation accumulation in selfers under different models of selection (rather than to contrast selfers and outcrossers). We find that genomes containing a larger proportion of loci under fluctuating selection experience higher fixation rates genome wide, both because of higher fixation rates at the fluctuating loci themselves but also through a linked selection effect on other loci, including those under constant selection. These results imply that loci under fluctuating selection contribute disproportionately to mutation accumulation and fitness decline.

## Methods

In most respects, the simulation follows standard assumptions so we only describe the important features here but provide a detailed description in Supplementary File Part 2 (simulation code available from https://github.com/EddieKHHo/SelfingMutAccum). The population consists of *N* diploid hermaphrodites. Each genome consists of 5000 bi-allelic loci that affect fitness as well as three neutral quantitative loci that are used to estimate effective population size, *N_e_*. Loci are arranged on a single linear chromosome that experiences, on average, *M* = 1 recombination event per gamete. Fitness acts multiplicatively across loci and we assume deleterious alleles are partially recessive, *h =* ¼. Reproduction occurs following standard assumptions assuming a sporophitic selfing of *S* = 0.98. Each offspring receives, on average, *U* new deleterious mutations. We confirmed the program produces the expected population genetic results under constant selection (detailed in Supplementary File Part 3).

## Base model

Our “base model” assumes individuals carry two types of loci that affect fitness. “C-loci” experience a constant selection coefficient of *s_c_* in all generations. “F-loci” experience a strength of selection that changes over time, *s_f_*(*t*), taking the value 0 or *s_max_*, i.e., mutations at F-loci are conditionally neutral. A proportion, *p_f_*, of the fitness affecting loci in the genome are F-loci and the remainder are C-loci; location assignment of locus types within the genome was randomized every simulation run. All F-loci experience the same selection in a given generation (0 or *s_max_*) as if they all are affected by the same environmental factor. The parameter *f* controls the temporal autocorrelation in selection. With probability *f* selection remains the same as the previous generation. With probability, 1-*f, s_f_*(*t*) is randomly chosen to be 0 or *s_max_* with probability 1- *ϕ* and *ϕ*, respectively; *ϕ* is the expected proportion of generations that *s_f_*(*t*) = *s_max_*. The geometric average strength of selection experienced by loci under fluctuating selection is 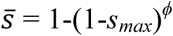. For all simulations we chose *s_max_* such that 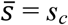, so that F-loci are expected to experience the same geometric average selection strength as C-loci to provide the fairest comparison between these types of loci.

After a burn-in period, we calculated the per locus fixation rates for C- and F-loci separately as the number of fixations at each type of loci divided by the number of generations over which the data was collected and divided by the number of loci for each type. Throughout, we calculated the *relative fixation rate* as the fixation rate in populations with *p_f_* > 0 divided by the fixation rate in populations with *p_f_* = 0 (holding all other parameters equal). The relative fixation rate for C-loci quantifies the extent to which fixation rates are affected by residing in genomes that experience fluctuating selection rather than residing in genomes that only experience constant selection (i.e., the classic model). The relative fixation rate for F-loci quantifies the direct effect of fluctuating selection as well as effect of residing in a genome experiencing fluctuating selection.

## Simulations with exponentially distributed selection

We modified the base model by allowing selection strength to vary over time following an exponential distribution (as opposed to the discrete “on/off’ version of conditional neutrality assumed in the base model). Each generation, consecutive environments remain the same with probability *f*. With probability, 1-*f*, the selection strength is chosen from an exponential distribution with mean 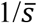. Similar to the base model, C- and F-loci experienced the 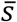. We focused on simulations with 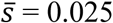 where the geometric mean and the variance in selection strength under the exponential model was almost identical to that of the conditionally neutral mutations with 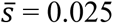 and *ϕ* = 0.5 in the base model.

## Simulations with two fluctuating ecological variables

An unrealistic assumption of the base model is that all F-loci experience fluctuating selection in exactly the same way. As the simplest way to relax this assumption, we allowed for two sets (F1, F2) of loci under fluctuating selection and we imagine that each set responds to a different ecological variable. These sets comprise *p_f1_* and *p_f2_* of the genome and the remaining proportion, *1-p_f1_-p_f2_*, are C-loci. We assumed equally sized sets of fluctuating loci*, p_f1_* = *p_f2_* (and *p_f1_* + *p_f2_* = *p_f_*).

If mutations at F-loci are assumed to be conditionally neutral (as in the base model), both F-loci sets experience fluctuations in selection between 0 and *s_max_* with the same 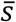. Each generation, the environment remains in the same state with probability *f*. With probability, 1-*f*, the environment is randomly selected among four possible states: (*s_f1_*(*t*) = *s_max_, s_f2_*(*t*) = *s_max_*}, {*s_f1_*(*t*) = *s_max_, s_f2_*(*t*) = 0}, (*s_f1_*(*t*) = 0, *s_f2_*(*t*) = *s_max_*}, (*s_f1_*(*t*) = 0, *s_f2_*(*t*) = 0} with probabilities *ϕ*^2^*+d, ϕ*(1-*ϕ*)−*d*, (1-*ϕ*)*ϕ*−*d*, (1-*ϕ*)(1-*ϕ*)+*d*, respectively. The value of *d* determines correlation *ρ* in selective states between sets F1 and F2. We also examined simulations with sets F1 and F2 in which selection fluctuates over time following an exponential distribution; selection between sets F1 and F2 was made to be either completely correlated, independent or negatively correlated (*ρ* = 1, 0, −0.645, respectively; detailed in Supplementary file Part 2).

## Simulations with beneficial mutations

Lastly, we extended the base model by allowing beneficial mutations to occur at C-loci with a genome wide rate of *U_b_*, assuming *U_b_* << *U*. We estimated the fixation probability for a new beneficial mutation as the number of beneficial fixations divided by the expected number of beneficials that were introduced (i.e., *N* * *U_b_* * number of generations).

### Results

#### Heuristic model

We first outline predictions for the fixation probability of a new deleterious mutation occurring at a locus experiencing temporal fluctuations in the intensity of negative selection, denoted as *P_f_*. These predictions are useful for interpreting the results of our multi-locus simulations. If selection fluctuates temporally between two selection coefficients *s=* {0, *s_max_*} with equal frequency (*ϕ* = 0.5), we can make simple predictions for *P_f_* in the limiting cases where selection fluctuates very often or very rarely. In the first case, selection fluctuates so often that a new mutation will experience many periods of neutrality and strong selection before fixation and thus experiences its expected (geometric) average fitness. We predict that *P_f_* would be similar to the fixation probability under time constant selection with 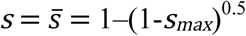; we denote this prediction as 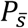. In the second case, selection fluctuates so rarely that a new mutation is almost certain to be lost or fixed before selection changes. The expected fixation probability is then 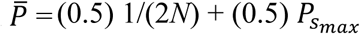, where 1/(2*N*) is the fixation probability for a neutral mutation (*s =* 0) and *P_Smax_* is the fixation probability when *s=s_max_*.

Given these two limiting cases, we make two predictions regarding *P_f_*. Because the fixation probability for a new deleterious mutation is a convex function of *s* (Fig. S1a; Kimura 1962), it follows that 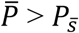 (via Jensen’s inequality) and we predict *P_f_* to lie within the boundaries represented by the limiting cases of 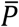 and 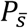 (i.e., 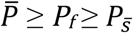; Fig. S1b, c). Second, as 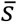 increases, 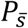 decays quickly and asymptotes at 0 while 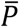 decays more slowly and asymptotes at 1/(4*N*) (Fig. S1b) so there is a larger proportional change in 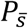 than 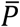 with respect to changes in 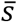 (Fig. S1e). This means that 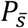 is more sensitive to changes in *s_max_* than 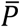; we predict *P_f_ to* be intermediate in its sensitivity to changes in 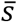 (Fig. S1b, e). We see a similar pattern when changing the population size, *N*. If we hold 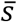 constant, we observe that 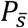 is more sensitive to changes in *N* compared to *P_f_* and 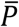 (Fig. S1c, d, f).

The heuristic arguments above are based on single locus theory. Below we investigate mutation accumulation in multi-locus models of a highly selfing species where selective interference plays an important role in elevating fixation rates; nonetheless, we show that the main ideas of the heuristic model apply. Within the context of our multi-locus simulations, comparing the deleterious fixation rates of loci experiencing constant selection (C-loci) and loci experiencing fluctuation selection (F-loci) is similar to comparing 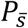 and *P_f_*, respectively (assuming both loci experience the same 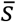). Thus, we predict fixation rates at F-loci to be higher than that of C-loci (because 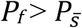). We also predict fixation rates at F-loci to be less sensitive to changes in 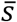 and *N_e_* than fixation rates at C-loci.

#### Temporal autocorrelaition in selection

Our heuristic model implies that the fixation probability of mutations should depend on the temporal autocorrelation of selection, *f* 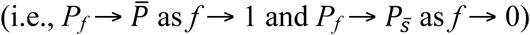. We confirmed this in our multi-locus model of highly selfing populations (Fig. S3, S4); Wardlaw and Agrawal (2012) found similar results for the deleterious fixation rate in obligately asexual populations. Rather than further exploring the effects of *f* on fixation rates, we assume that selection undergoes moderate degrees of temporal autocorrelation (*f* = 0.95, *ϕ* = 0.5) such that the correlation in selection between time points separated by 1, 10, and 100 generations is 0.95, 0.60, and 0.006, and the average run length between switches in the environment is ~40 generations. This regime elevates fixation rates at loci under fluctuating loci, but is moderate in its effects; fixation rates can be much higher with higher values of *f* and lower values of *ϕ* (Fig. S3, S4, S15). In the remaining sections, we focus on exploring how loci under fluctuating selection in the genome alter fixation rates at linked sites and genome wide.

#### Varying the composition of the genome, p_f_

We examine fixation rates in genomes where a fraction *p_f_* of loci are under fluctuating selection (F-loci) and the remaining loci are under constant selection (C-loci). Unless otherwise noted, we focus on simulations of highly selfing populations (*S* = 0.98) with *N* = 10^4^, 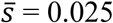, and *h* = 0.25 with *f* = 0.95 and *ϕ* = 0.5; results for obligately selfing populations (*S* = 1) can be found in Supplementary File Part 1. Per locus fixation rates within genomes experiencing fluctuating selection (*p_f_* > 0) are typically higher than fixation rates in the classic model (*p_f_* = 0); this is true for both C- and F-loci, but the effect is stronger for the latter (Fig. 1) The fact that per locus fixation rates vary with *p_f_* indicates that changing the density of F-loci in the genome alters the effects of linked selection on both F- and C-loci. When less than half the genome consists of F-loci (*p_f_* < 0.5), increasing *p_f_* generally elevates fixation rates at both C- and F-loci and, thereby, the genome-wide average fixation rate. At high mutation rates (*U* = 1) further increasing *p_f_* reduces fixation rates for genomes that contain more than 50% F-loci (Fig. 1a). As it seems unlikely that a high fraction of loci experience strong fluctuations in selection, the results for *p_f_* < 0.5 are the most biologically relevant.

**Figure 1.**
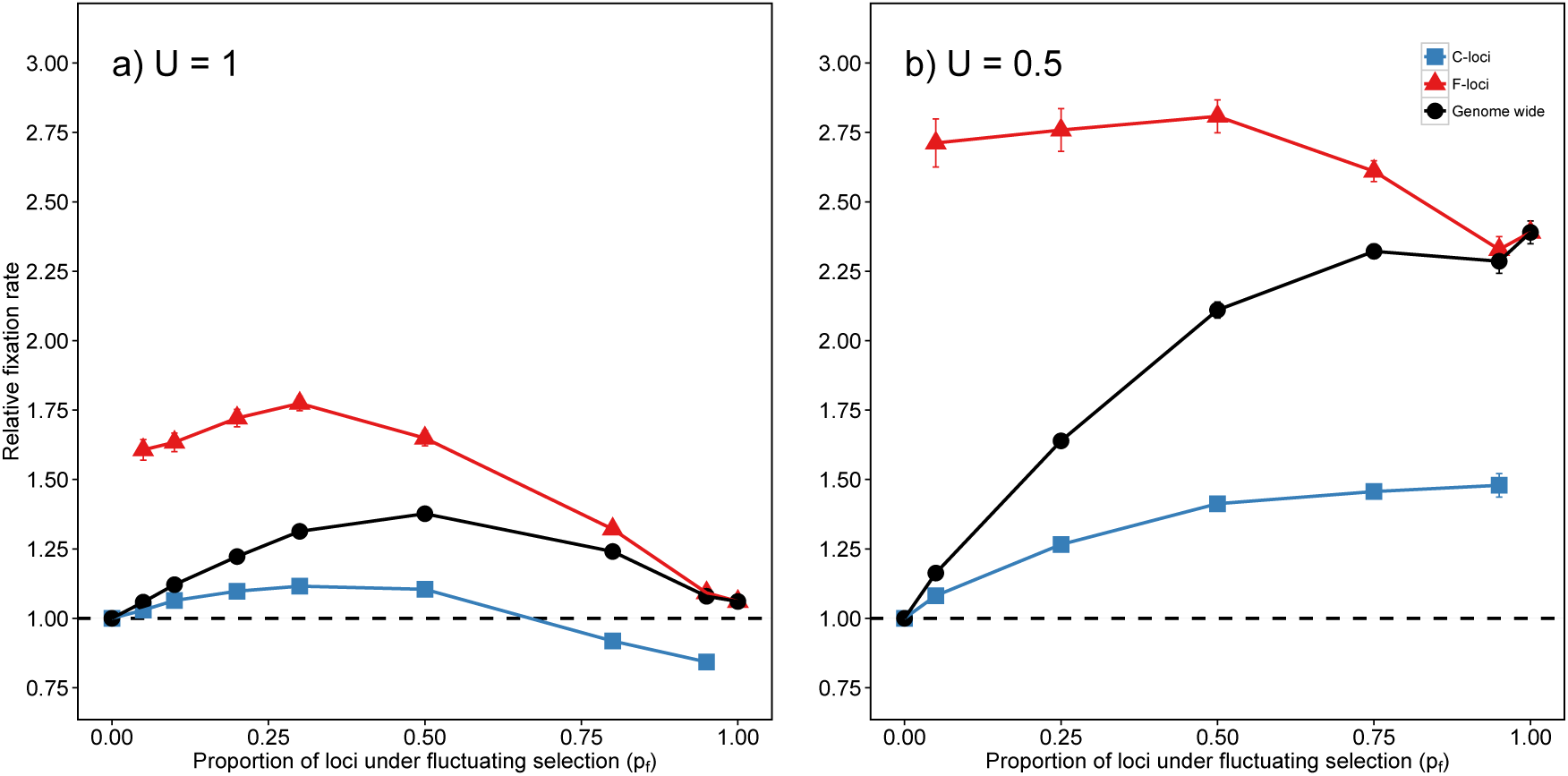
Relative fixation rates (mean ± SE) for C- (blue) and F-loci (red) in highly selfing populations with varying genomic composition, *p_f_*. Fixation rates under high mutation rates in (a) ***U*** = 1 and low mutation rates (b) ***U*** = 0.5. Fixation rates are given relative to the fixation rate for C-loci in the classic model (*p_f_* = 0), which is represented by the horizontal dashed line at 1.0 (the grey shading denotes SE). The absolute fixation rates in the classic model (*p_f_* = 0) for panels a,b are 7.47 × 10^−6^ and 8.74 × 10^−7^ per locus per generation respectively. Other parameters: *N* = 10^4^, *S* = 0.98, *f* = 0.95, *ϕ* = 0.5, *h* = 0.25, 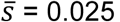.

When mutation rates are lower (so absolute fixation rates are lower), the effect of fluctuating selection on the *relative* fixation rate is more dramatic for both C- and F-loci. For the parameter sets examined in Fig. 1, the inclusion of F-loci maximally increases genome-wide fixation rate by a factor of 1.38 vs. 2.42 at the high and low mutation rate, respectively. This effect of mutation rate holds in simulations using other values of autocorrelation *f* and frequency of selection *ϕ* and in obligately selfing populations (Fig. S4, S5). Although the relative impact of fluctuating selection is larger at lower mutation rates, in the remaining sections we focus on results from high mutation rates (*U* = 1) because simulations at lower mutation rates require much longer runs to obtain accurate estimates of fixation rates.

The density of F-loci alters the effective population size, *N_e_* (Fig. 2a shows *N_e_* for the simulations represented by Fig. 1; Fig. S6 for obligately selfing populations). This might be the reason for the variation in fixation rates in relation to *p_f_* (Fig. 1a). If so, fixation rates should be equal for simulations with different values of *p_f_* if they experienced the same *N_e_*. To test this, we ran additional simulations changing the census population size, *N*, of the simulations to raise or lower *N_e_* for a given value of *p_f_*. (We observe that *N_e_* is not linearly related to *N* (Fig. 2b) as has been observed previously in simulations of asexual populations accumulating deleterious mutations due to Muller’s ratchet (Gordo et al. 2002).) Consistent with the idea that the relationship between *p_f_* and fixation rate results from the effect of *p_f_* on *N_e_*, fixation rates are similar across different values of *p_f_* when *N_e_* is similar, but only if *p_f_* is not too high (0 < *p_f_ ≤* 0.5; Figs. 2c, d). However, when comparing simulations with similar estimates of *N_e_*, very high values of *p_f_* (i.e., *p_f_* = 0.8, 0.95) result in higher rates fixation rates than low or moderate values of *p_f_* (i.e., *p_f_ ≤* 0.5). This indicates that when *p_f_* is large (and the mutation rate is high), selective interference between C- and F-loci affects fixation rates in ways that cannot be captured by a genome-wide estimate of *N_e_*.

**Figure 2.**
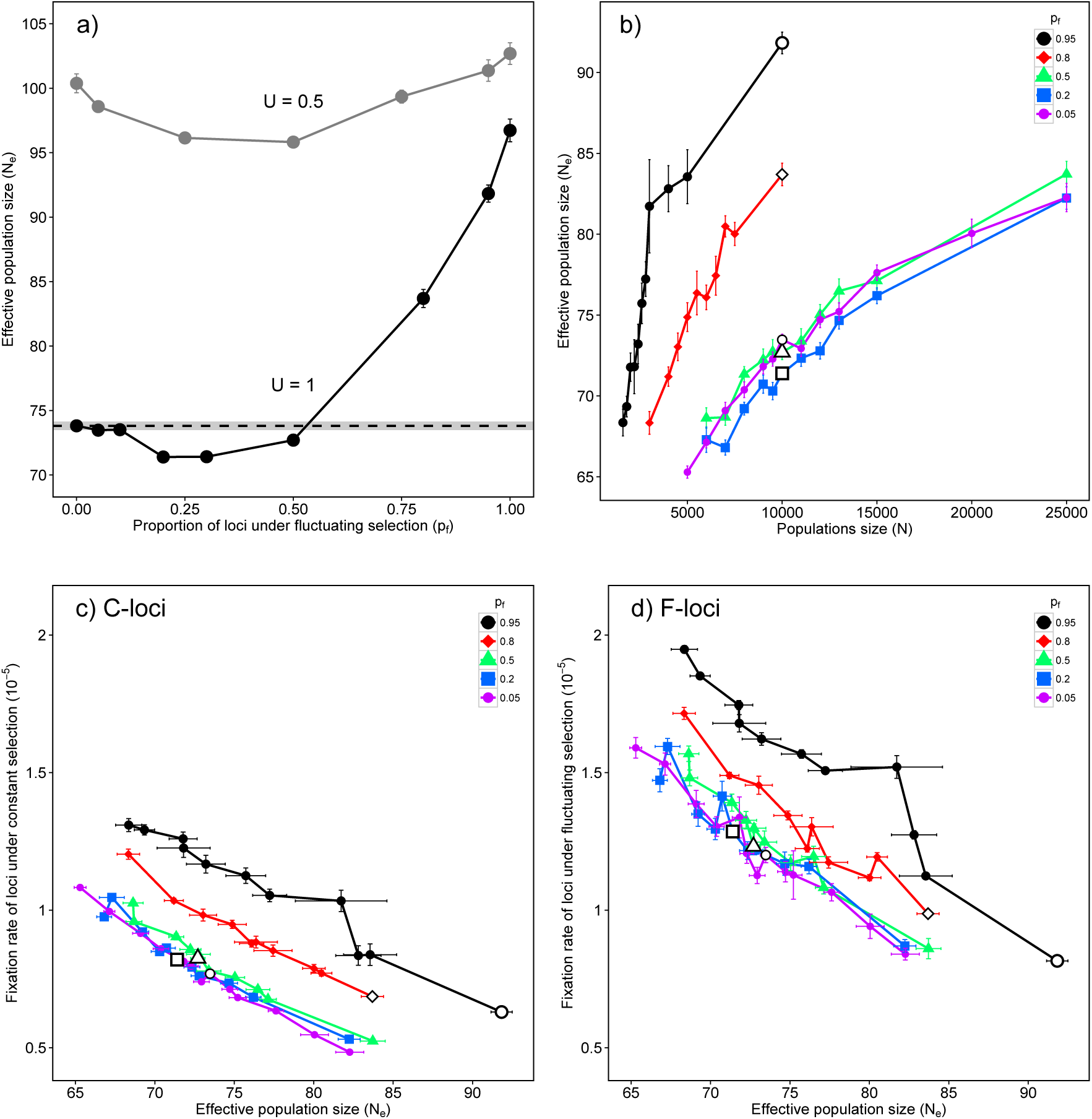
(a) Estimates of effective population size (mean ± SE) for genomes containing different densities of F-loci, *p_f_*, in a population of *N* = 10^4^. Shown are *N_e_* values for highly selfing (*S* = 0.98) populations under high (*U* = 1) and lower (*U* = 0.5) mutation rates; these results are from the same simulations represented in Figs. 1a,b. The horizontal dashed line indicates the *N_e_* (grey shading is SE) for the corresponding classic model (*p_f_* = 0). (b) Estimates of *N_e_* plotted against *N* (*U* = 1) for genomes with different values of *p_f_* as depicted in the legend. The open symbol indicates the *N_e_* when *N* = 10^4^ (i.e., corresponding to simulations in Fig. 1a). Fixation rates plotted against *N_e_* (c) loci under constant selection (d) loci under fluctuating selection (*U* = 1).

#### Fixation bias at F-loci

As predicted from the heuristic model, the fixation rate of deleterious mutations at F-loci is always greater than that for C-loci within all mixed genomes (Figs. 1, 2c, d). This bias is clearly stronger when selection fluctuates in ways that directly increase fixation rates at F-loci (i.e., larger *f* and smaller *ϕ* Figs. 3a, S4). Moreover, this bias is affected by the ‘traditional’ parameters affecting mutation accumulation such as the strength of selection 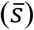, the mutation rate (*U*) and the population size (*N*). To examine how this bias depends on aspects of selection, we performed simulations where half the genome consist of F-loci (*p_f_* = 0.5) with an average selection strength of 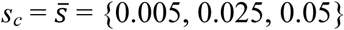. The bias in fixations at F-loci increases with 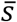 as stronger selection reduces fixation rates at C-loci relatively more than at F-loci (Fig. 3b). These results can be interpreted in light of the heuristic model that predicts F-loci will be less sensitive than C-loci to increases in 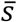: mutations at F-loci can accumulate during episodes where selection is weak regardless of how large *s_max_* becomes in episodes when selection is strong. Overall, conditions that slow down mutation accumulation, such as stronger 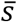 (Fig. 3), lower *U* (Figs. 4, S4, S14b), larger *N* (Fig. S9), and less selfing (Fig. S10) reduce fixation rates at C-loci more so than at F-loci, increasing the relative contribution of F-loci to total mutation accumulation; see Fig. S8 for simulations of obligately selfing populations.

**Figure 3.**
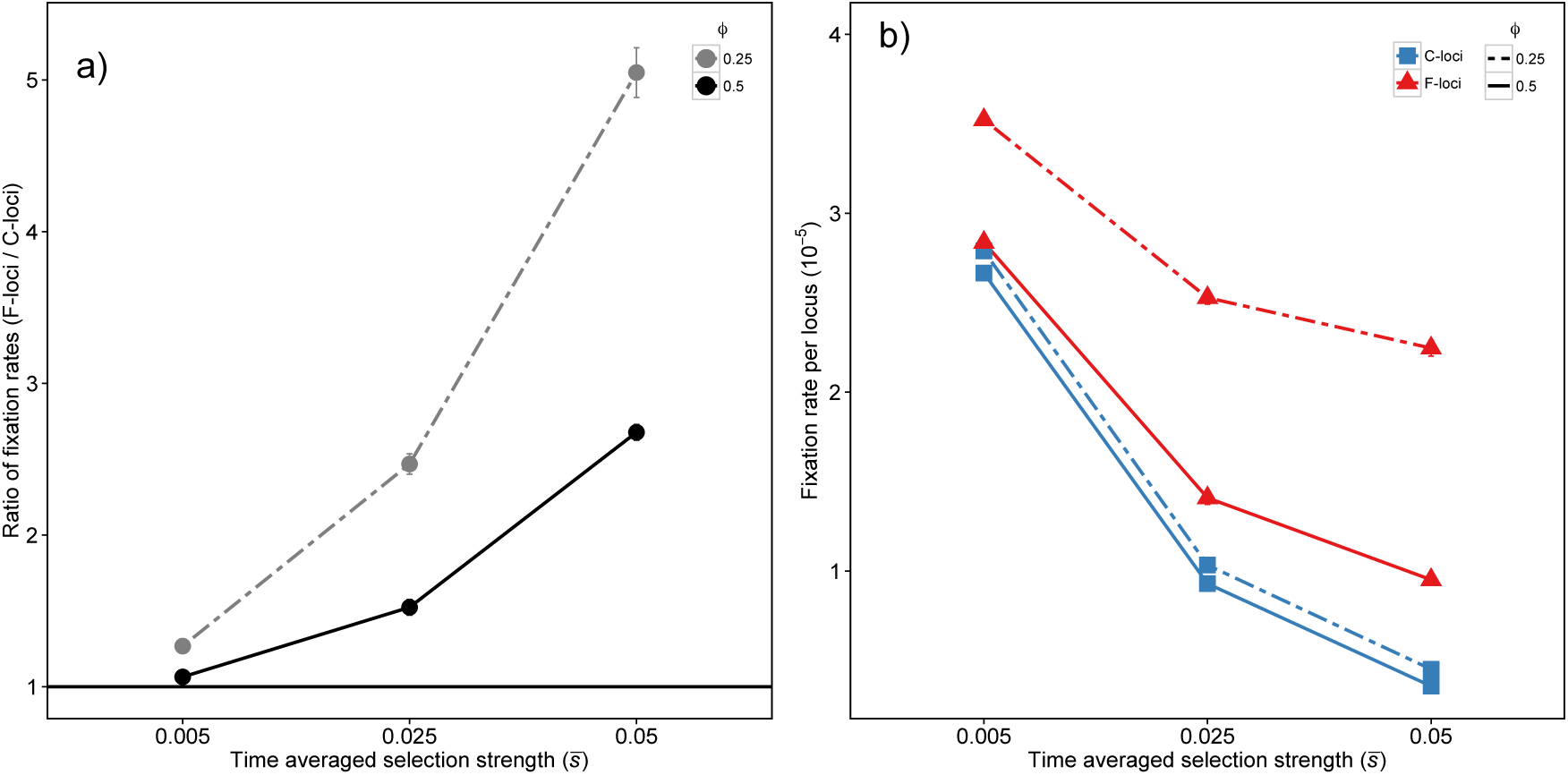
(a) Ratio of fixation rates at F-loci to fixation rates at C-loci when half of the genome consists of F-loci (*p_f_* = 0.5); error bars are SE (often too small to be visible). When ***ϕ*** = 0.5 (solid black line), selection is non-neutral (*s_max_*) half the time but when ***ϕ*** = 0.25 (double-dashed grey line), selection is non-neutral only a quarter of the time. To maintain 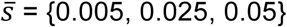 when ***ϕ*** is reduced by half we approximately double the value of *s_max_* (Methods), therefore making selection concentrated into shorter but more intense events. (b) Fixation rates at F-loci (red) and C-loci (blue) when ***ϕ*** = 0.5 (solid line) and when ***ϕ*** = 0.25 (double-dashed line). Other parameters: *S* = 0.98, *N* = 10^4^, *U* = 1, *h* = 0.25, *f* = 0.95.

**Figure 4.**
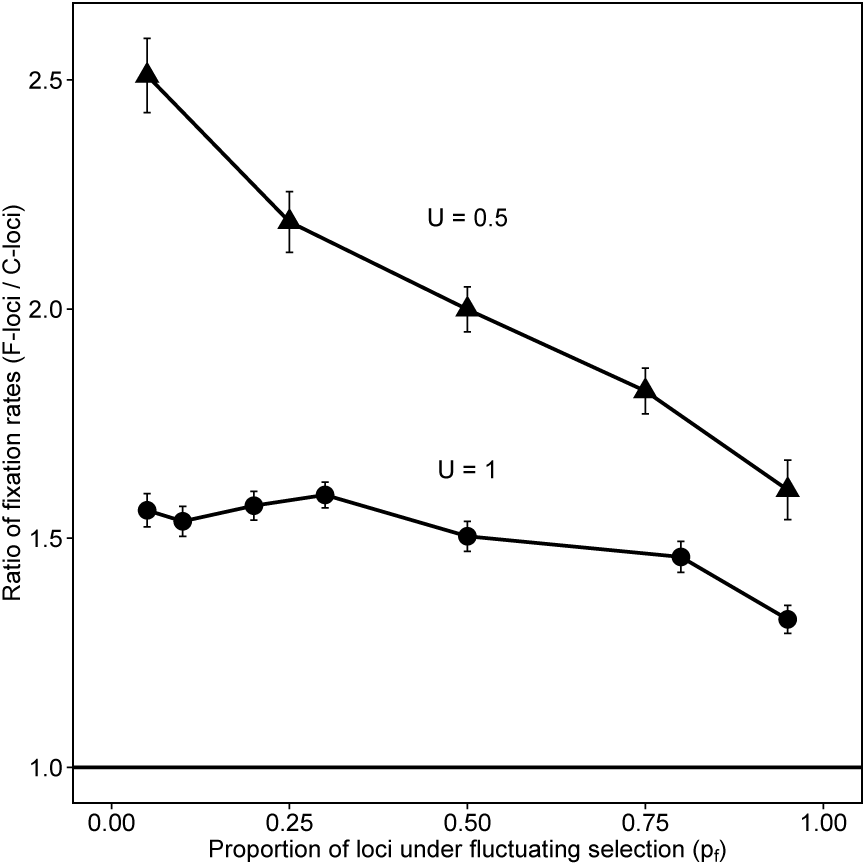
Ratio of fixation rates (mean ± SE) at F-loci to fixation rates at C-loci in populations with different proportions of F-loci under high (*U* = 1, circles) and low (*U* = 0.5, triangles) mutation rates. Other parameters: *S* = 0.98, *N* = 10^4^, *h* = 0.25, *f* = 0.95, *ϕ* = 0.5.

#### Exponentially distributed selection over time

In the base model we assumed that mutations at F-loci are conditionally neutral with selection fluctuating discretely between 0 and *s_max_*. As a test of whether the patterns above are robust to that assumption, we consider simulations where the selection strength at F-loci is randomly sampled from an exponential distribution when the environment changes. Both models have a similar mean and standard deviation in selection strength (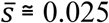, standard deviation in *s_f_*(*t*) = 0.025). Using *f* = 0.95 in the exponential model (as in the base model), the correlation in selection between time points separated by 1, 10, and 100 generations is 0.95, 0.60, and ~0, respectively. The fixation rates for both C- and F-loci are slightly lower under the exponential model than the discrete conditional neutrality model (Figs. 5, S12). Mutations fix disproportionately at F-loci under the exponential selection model, but this bias is not quite as strong as observed in the conditional neutrality model. As in the base model, reducing *U* (from *U* = 1 to *U* = 0.5) under the exponential model increases relative fixation rates and the proportion of mutations that fix at F-loci (Figs. 5, S14). Though the base model and exponential model represent qualitatively different forms of temporal heterogeneity in selection, the effects of these models on fixation rates are reasonably similar. We speculate that any form of fluctuations in selection intensity are likely to elevate fixation rates and fitness decline in highly selfing populations as long as the variance in selection over time is sufficiently large and there is some degree of temporal autocorrelation.

**Figure 5.**
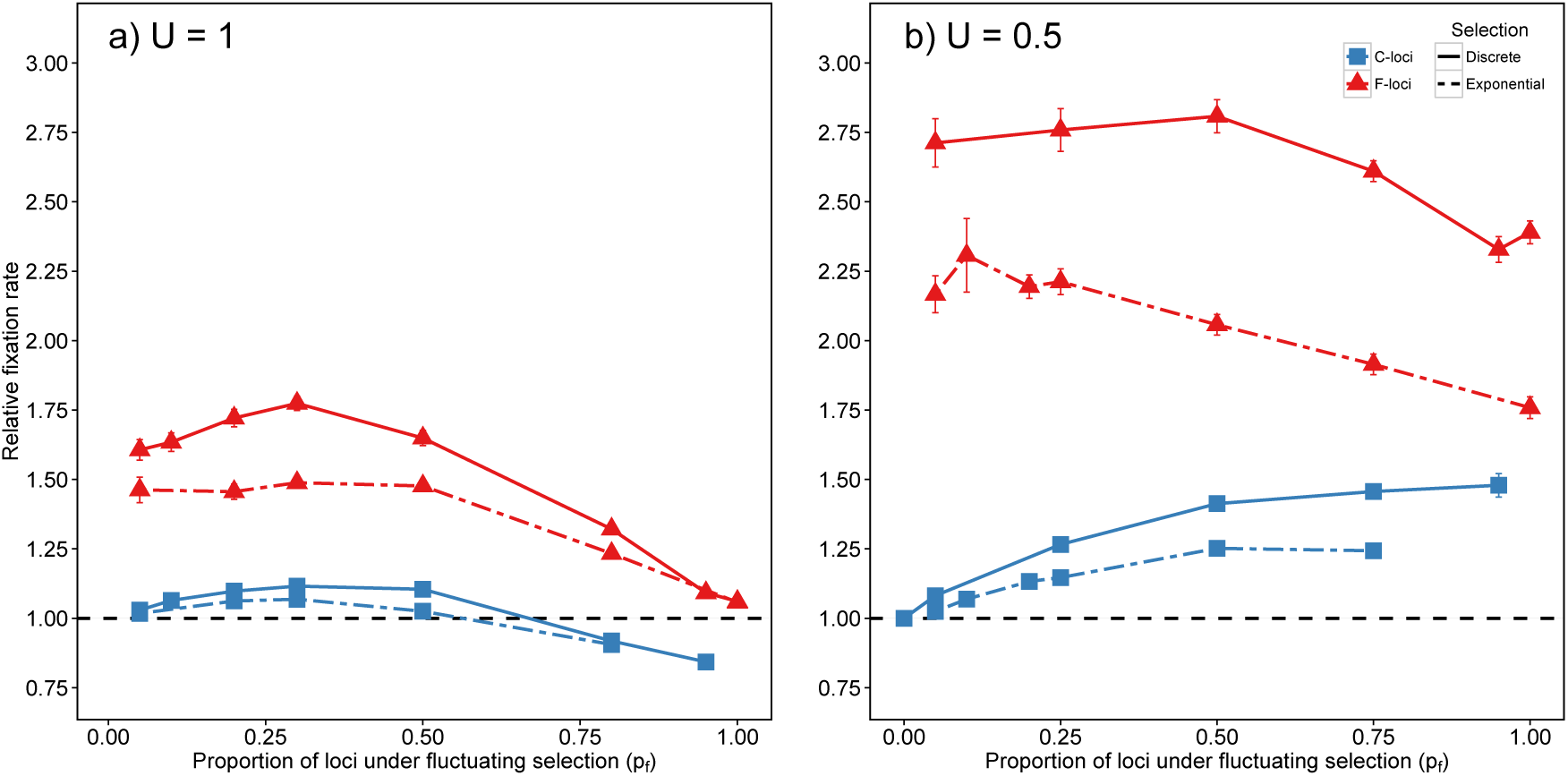
Relative fixation rates (mean ± SE) at C- (blue) and F-loci (red) in populations with different genomic compositions, *p_f_*, when selection strength is modelled as discrete conditional neutrality (solid line) or following an exponential distribution (double-dashed line) over time under (a) high mutation rates (*U* = 1) and (b) low mutation rates (*U* = 0.5). Relative fixation rates under the conditional neutrality model are re-plotted from Figs. 1a,b. The geometric time average and the variance in selection strength and for the conditional neutrality and exponential selection model are approximately the same (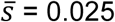, standard deviation in *s_f_*(*t*) = 0.025). Fixation rates are given relative to the fixation rate for C-loci in the classic model (*p_f_* = 0), which is represented by the horizontal dashed line at 1.0 (the grey shading denotes SE). Other parameters: *S* = 0.98, *f* = 0.95, *N* = 10^4^, *h* = 0.25.

#### Two components of fluctuating selection

In the previous sections, we assumed that all F-loci simultaneously respond to the same environmental factor that fluctuates in time. As the simplest way to relax this assumption, we allow for two equally sized sets of F-loci (F1, F2) that respond to two separate environmental factors. The two environmental factors may fluctuate in perfect synchrony, causing a complete correlation in selective states between loci within F1 and F2 (*ρ* = 1), which is equivalent to the base model. Alternatively, the two environment factors fluctuate in anti-synchrony, causing a negative correlation in selective states between F1 and F2 (*ρ* = −1). We perform simulations for a range of different values of *ρ*, the correlation in selective states between set F1 and F2.

The environmental correlation *ρ* has little effect on the fixation rates of C- and F-loci when the genome consists of mostly C-loci (*p_f_ ≤* 0.5); *ρ* is negatively related to fixation rate when the genome consists of mostly F-loci, making our base model (*ρ* = 1) conservative in this respect (Fig. 6a); obligately selfing populations are less insensitive to changes in *ρ* (Fig. S12). We observe qualitatively similar effects of *ρ* when selection between the two groups is exponentially distributed in time (Figs. 6b, S12, S13). In sum, under biologically realistic ranges of *p_f_* (i.e., *p_f_* < 0.5), fixation rates do not appear to be strongly dependent on how fluctuations in selection are correlated between two equally sized sets of F-loci.

**Figure 6.**
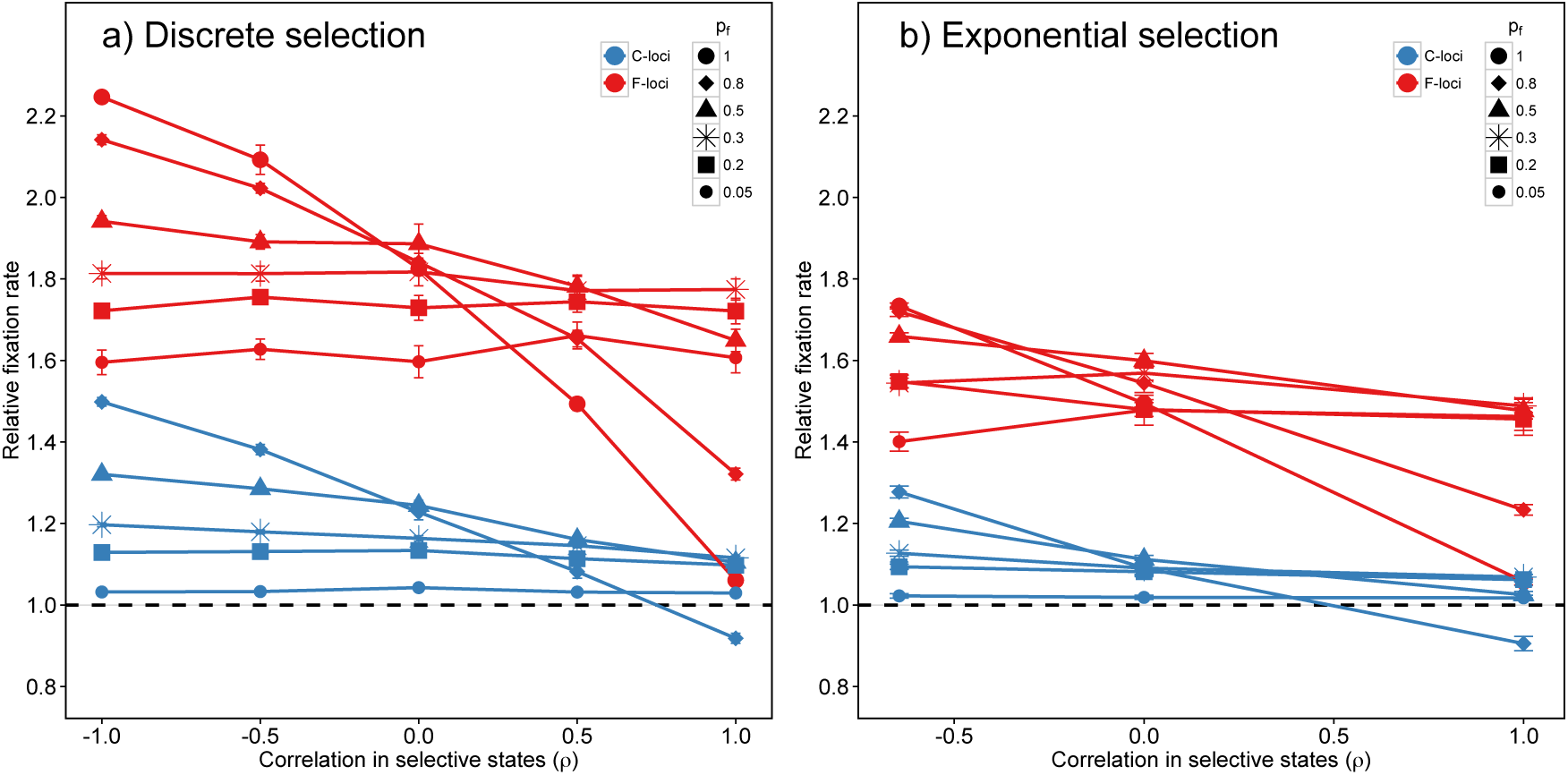
Relative fixation rates (mean ± SE) for C-loci (blue) and F-loci (red) as a function of the correlation in environmental variables responsible for fluctuating selection *ρ*. Loci under fluctuating selection are equally distributed between two sets of F-loci (F1, F2). The correlation in selective states between F1 and F2, *ρ*, can be positive or negative or zero. The F-loci (summing both F1 and F2) comprise a fraction *p_f_* of the genome. (a) Selection under the model of discrete conditional neutrality; *ρ* = {1, 0.5, 0, −0.5, 1}. (b) Selection is exponentially distributed over time; *ρ* = {1, 0, −0.645}. Fixation rates are shown for genomes with different frequencies of fluctuating loci, *p_f_*. Fixation rates are given relative to the fixation rate for C-loci in the classic model (*p_f_* = 0), which is represented by the horizontal dashed line at 1.0 (the grey shading denotes SE); the actual fixation rates is 7.47 × 10^−6^. Other parameters: *S* = 0.98, *f* = 0.95, ***ϕ*** = 0.5, 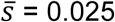, *N* = 10^4^, *U* = 1, *h* = 0.25.

#### Beneficial mutations

We performed simulations in which beneficial mutations occurred at C-loci at much lower rates than deleterious mutations (*U_b_ ≤* 0.001) and had smaller, larger or equal fitness effects compared to deleterious mutations (beneficials: *s_b_* = (0.01, 0.025, 0.05, 0.08}, *h_b_* = 0.5; deleterious: 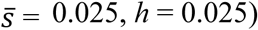, *h* = 0.025). F-loci only received deleterious mutations and experienced selection fluctuating discretely over time between 0 and *s_max_* (*f* = 0.95, *ϕ* = 0.5).

The fixation probability of beneficial mutations was lower in simulations with F-loci (*p_f_*= 0.2) than those without F-loci (*p_f_* = 0) for all values of *s_b_* (Table 1) indicating that deleterious mutations at F-loci caused more interference with beneficials than those at C-loci. Beneficials with smaller or similar effect size to deleterious mutations (i.e. *s_b_* = 0.01, 0.025) had little effect on the deleterious fixation rate. However, beneficials with larger fitness effects (*s_b_* = 0.05, 0.08) increased deleterious fixation rates at both C- and F-loci (Table 1). The deleterious fixation rate of C-loci changed proportionately more with increases in the strength of beneficials (*s_b_*) compared to that of F-loci (Table 1), consistent with our heuristic model. Recall that comparing fixation rates at F-loci and C-loci is similar to comparing *P_f_* and 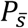, respectively, and 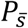 is proportionally more sensitive to changes in *N* and *s* compared to *P_f_* (Fig. S1e, f). Because an increase in *s_b_* causes a decrease in *N_e_* (Table 1), we expect C-loci 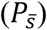 to be more sensitive to this change than F-loci (*P_f_*).

Because the F-locus rate without beneficials is larger than the C-locus rate, the fact that the absolute increase in the deleterious fixation rate caused by beneficials is larger for F-loci than C-loci is obscured by only looking at their “proportional increase”. This is clearer by taking a second perspective in which we calculate the average number of additional fixed deleterious mutations per fixed beneficial (Table S3). When the beneficial effect size is larger than deleterious (*s_b_* = 0.05, 0.08), each beneficial that fixes is responsible for, on average, less than one additional deleterious fixation but these are not evenly distributed between F- and C-loci. F- loci are only 25% as abundant as C-loci (i.e., *p_f_* = 0.2), but the number of additional fixed deleterious F-alleles per fixed beneficial is more than 25% the number of additional fixed deleterious C-alleles. From this perspective, F-loci experience stronger interference from beneficials, in an absolute sense, than do C-loci even though C-loci experience a greater proportional interference effect than F-loci (Table 1).

**Table 1.** Summary of simulations with beneficial mutations

| $U$ | $U_b$ | $s_b$ | $p_f$ | $N_e$ (SE) | Ben prob fix<br>x $10^{-3}$ (SE) | Del fix rate relative to NO ben (SE) | |
| --- | --- | --- | --- | --- | --- | --- | --- |
|  |  |  |  |  |  | C-loci | F-loci |
| 0.5 | 0.0001 | 0.01 | 0 | 99.7 | 0.172 | 0.992 |  |
|  |  |  |  | (0.35) | (0.004) | (0.006) |  |
|  |  |  | 0.2 | 96.5 | 0.164 | 0.999 | 1.032 |
|  |  |  |  | (0.36) | (0.005) | (0.005) | (0.013) |
|  |  | 0.025 | 0 | 99.7 | 0.752 | 1.021 |  |
|  |  |  |  | (0.37) | (0.011) | (0.009) |  |
|  |  |  | 0.2 | 96.9 | 0.703 | 1.019 | 1.072 |
|  |  |  |  | (0.48) | (0.012) | (0.007) | (0.018) |
|  |  | 0.05 | 0 | 96.6 | 3.798 | 1.312 |  |
|  |  |  |  | (0.25) | (0.015) | (0.008) |  |
|  |  |  | 0.2 | 93.5 | 3.499 | 1.240 | 1.159 |
|  |  |  |  | (0.33) | (0.025) | (0.008) | (0.024) |
|  |  | 0.08 | 0 | 87.5 | 11.429 | 2.373 |  |
|  |  |  |  | (0.34) | (0.040) | (0.011) |  |
|  |  |  | 0.2 | 85.9 | 10.631 | 2.092 | 1.664 |
|  |  |  |  | (0.34) | (0.044) | (0.012) | (0.022) |
| 1 | 0.001 | 0.01 | 0 | 73.8 | 0.121 | 0.994 |  |
|  |  |  |  | (0.33) | (0.002) | (0.003) |  |
|  |  |  | 0.2 | 71.2 | 0.119 | 0.980 | 1.039 |
|  |  |  |  | (0.24) | (0.002) | (0.003) | (0.018) |
|  |  | 0.025 | 0 | 72.8 | 0.384 | 1.016 |  |
|  |  |  |  | (0.34) | (0.003) | (0.003) |  |
|  |  |  | 0.2 | 71.0 | 0.356 | 1.005 | 1.008 |
|  |  |  |  | (0.33) | (0.004) | (0.003) | (0.011) |
|  |  | 0.05 | 0 | 71.0 | 1.531 | 1.199 |  |
|  |  |  |  | (0.31) | (0.007) | (0.004) |  |
|  |  |  | 0.2 | 67.7 | 1.405 | 1.152 | 1.129 |
|  |  |  |  | (0.31) | (0.007) | (0.004) | (0.018) |
|  |  | 0.08 | 0 | 62.5 | 3.916 | 1.774 |  |
|  |  |  |  | (0.29) | (0.010) | (0.003) |  |
|  |  |  | 0.2 | 60.9 | 3.672 | 1.650 | 1.478 |
|  |  |  |  | (0.28) | (0.008) | (0.004) | (0.012) |

## Discussion

Because of the reduced efficacy of selection associated with high selfing rates, such populations can accumulate and permanently fix deleterious mutations at high rates leading to rapid fitness decline and possibly to extinction (Heller and Maynard Smith 1979, Charlesworth and Charlesworth 1993). Environmental changes occur on many different times scales (Halley 1996, Vasseur and Yodviz 2004, Furguson et al. 2016) and can induce fluctuations in the intensity and direction of selection (Siepielski et al. 2017, Bell 2010) but the consequences for mutation accumulation in selfers is unstudied. Loci experiencing selection that fluctuates in intensity, F-loci, fix deleterious mutations at a higher rate than loci under constant selection, C-loci, experiencing the same time averaged selection strength 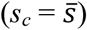. The extent to which F-loci contribute disproportionately to total mutation accumulation depends on aspects of environmental heterogeneity. The bias is stronger with increasing temporal autocorrelation in selection, *f*, and when selection is concentrated into rarer but more intense episodes (low values of *ϕ*), holding 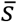 constant (Fig. S3, S4, S14). This bias also becomes stronger with changes to “traditional” parameters that lower the overall rate of mutation accumulation, i.e., higher values of 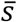 or *N* as well as lower values of *U* (Fig. 3, 4, S8, S9). In sum, fluctuating selection can increase the rate of mutation accumulation in highly selfing populations by several fold and the loci under fluctuating selection contribute disproportionately to mutational decline. These conclusions appear qualitatively robust to the exact nature of fluctuations (Fig. 5, 6, S12, S13).

In our model, deleterious mutations fix as a consequence of selective interference between linked sites. Supporting this, simulations with the same parameter values but at higher rates of outcrossing (*S ≤* 0.9) fix few to no mutations (Table S1). Obligately asexual organisms (Wardlaw and Agrawal 2012) and obligate selfers (*S* = 1) have high rates of mutation accumulation that are further increased by fluctuating selection. These effects remain important even with low levels of outcrossing (*S* = 0.98) that permit some genetic mixing. Interference between constantly selected loci can lead to the accumulation of deleterious mutations in asexual and highly selfing populations, a process classically called Muller’s ratchet (Haigh 1978, Heller and Maynard Smith 1979). However, little attention has been given to understanding of selective interference between loci under fluctuating selection and loci under constant selection.

In genomes containing a mix of F- and C-loci, F-loci have inherently higher rates of fixation but also tend to increase fixation rates at C-loci and other F-loci as illustrated in Fig. 1 where increasing the fraction of F-loci in the genome increases the per-locus fixation rates at both C- and F-loci (for 0 < *p_f_* < 0.5). This indicates that F-loci cause more selective interference (with both C- and F-loci) than C-loci. Similarly, we found that deleterious mutations at F-loci reduce fixation rates of beneficials more than deleterious mutations at C-loci do (Table 1, S2). Previous work in asexual populations where selection is constant over time but varies among loci shows that interference from strongly deleterious alleles increase fixation of more weakly deleterious alleles (Gordo and Charlesworth 2001, Soderberg and Berg 2007). From this we can infer that strong interference caused by F-loci occurs during periods when selection is strong, *s_f_*(*t*) = *s_max_* > *s_c_*. However, during periods where *s_f_*(*t*) = 0, F-loci do not cause interference. Given the observed net increase in fixation rate at C-loci, we infer that the increased interference when selection at F-loci is strong (*s_f_*(*t*) = *s_max_*) outweighs the reduced interference when selection is weak (*s_f_*(*t*) = 0). On the other hand, we observed that strongly selected beneficials interfered more strongly with deleterious alleles at C-loci than at F-loci in terms of the proportional change in their fixation rates (Table 1). Viewing interference from the beneficials as a reduction in *N_e_*, the result above is consistent with the heuristic model prediction that fixation probabilities for F- loci are less sensitive to *N_e_* than for C-loci.

Indeed the effect of selective interference on fixation rates can be interpreted as resulting from a reduction in the effective population size in some cases (Hill and Robertson 1966, McVean and Charlesworth 2000, Gordo and Charlesworth 2002). We observed that per locus fixation rates varied as a function of *p_f_* for both C- and F-loci (Fig. 1). In the range where *p_f_* was not too large (0 < *p_f_ ≤* 0.5), fixation rates for each locus type (F and C) were nearly equal for different values of *p_f_* when we adjusted *N* to obtain similar *N_e_* values (Fig. 2). Because neutral diversity (used to infer *N_e_*) and fixation rates are similarly affected by selective interference within this range of *p_f_* values, we conclude that selective interference in this regime can be thought of as causing a reduction in *N_e_*. Good et al. (2014) showed that in asexual populations experiencing selective interference the variance in fitness explains neutral diversity much better than the *N_e_* expected under deterministic mutation-selection balance. However, the time-averaged variance in fitness did not explain the variation in fixation rates in relation to *p_f_* better than our diversity-based estimates of *N_e_* (Fig. S7).

Neither *N_e_* nor the variance in fitness can account for the variation in fixation rates when most loci undergo fluctuating selection (*p_f_* > 0.5). Others have noted that the effects of selective interference on fixation rates cannot always be captured through *N_e_*. A sweep of a strongly beneficial mutation differentially affects neutral diversity and fixation probability at a linked weakly beneficial mutation (Stephan et al. 1992, Barton 1994). Fluctuating selection may have some sweep-like properties because genotypes favoured during the selective period (*s_f_*(*t*) = *s_max_*) can be different from those favoured during the neutral period (*s_f_*(*t*) = 0) due to the shift in the relative importance of F- and C-loci in determining fitness. Recurrent sweeps of highly fit genotypes caused by changing selective environments may represent another dynamical process that differentially affects fixation probability at selected loci and genetic drift at linked neutral loci (Comeron et al. 2008).

There are several questions in assessing the relevance of our model to natural systems. As highlighted in the Introduction, numerous lab studies show selection on mutations varies in strength across the environmental conditions. However, we do not know the extent to which this variation occurs in nature. Further, we have no estimate of the proportion of loci in the genome that experiences fluctuating selection, *p_f_*. Excluding mutations causing intrinsic lethality, we speculate that almost every locus experiences temporal change in selection strength as it seems implausible selection strength is exactly the same in all conditions. However, for many loci, fluctuations in selection may be minor. Perhaps only a minority of mutations undergo fluctuations in selection strength on the order of magnitude that we modeled and with at least a moderate level of temporal autocorrelation.

Environmental changes that cause fluctuations in selection may also cause fluctuations in population size (e.g., Bergland et al. 2014). Our main goal was to isolate the effects of fluctuating selection on its own but population size changes may be important on their own and/or interact with fluctuating selection. As described in Supplementary File Part 5, we briefly explored the interaction between fluctuating *N* and *s*. As expected, simulations with only fluctuating *N* fixed deleterious mutations at a rate similar to one with constant *N* = *N_h_*, where *N_h_* represents the harmonic time average population size (Otto and Whitlock 1997). Fluctuations in *s* alone increased deleterious fixation rates compared with one with constant 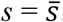. Keeping *N_h_* constant, the addition of fluctuating *N* into models of fluctuating s did not greatly change the deleterious fixation rate of the latter regardless of the (positive and negative) correlations in the fluctuations of *N* and *s*. In summary, fluctuations in *N* and *s* have different effects on deleterious fixation rates; we cannot simply use fluctuations in the product *Ns* as a substitute for fluctuations in each component alone.

Our study was motivated by the ‘dead end’ hypothesis for highly selfing species (Stebbins 1957, Igic and Busch 2013) but we do not explicitly model extinction. Similar to previous examinations of mutation accumulation in genomes with low rates of recombination (Haigh 1978, Heller and Maynard Smith 1979, Charlesworth and Charlesworth 1997), we hypothesize that deleterious fixations should lead to fitness loss and eventual extinction. Models using realistic parameters to predict the expected time to extinction under Muller’s ratchet assume deleterious mutations directly reduce population growth rates (Gabriel et al. 1993, Loewe 2006, Loewe and Lamatsch 2008). However, it is not obvious how the deleterious fitness effects of mutations directly affect ecological success and species persistence (Agrawal and Whitlock 2012). That said, our model predicts that mutations accumulate disproportionately at loci experiencing fluctuating selection. We might expect that populations are driven to extinction when deleterious mutations fix at loci required for surviving rare but severe climatic events. Prior to extinction, the central prediction is that F-loci contribute disproportionately to mutation accumulation. In reality, this would be challenging to test because it would be difficult to categorize the genome into F- and C-loci. Even if one did so (perhaps based on plasticity in expression), it would be extremely difficult to identify subsets of loci from each category that have comparable average selection strengths.

We focused on the mutation accumulation in highly selfing populations but the dynamics of deleterious mutations can affect the evolution of the selfing rate itself. In finite populations, the build up of negative linkage disequilibrium (LD) at sites under negative selection can increase deleterious fixation rates and favour the evolution of increased recombination and outcrossing (Keightley and Otto 2006, Gordos and Campos 2008, Kamran-Disfani and Agrawal 2014). Kamran-Disfani and Agrawal (2014) found that selfing rates evolve to just below where Muller’s ratchet begins turning at high rates. Because the presence of F-loci in the genome increases the deleterious fixation rate, it is natural to ask whether it also affects the evolution of selfing. In Supplementary File Part 4, we briefly explored this by allowing selfing rates to evolve in our model. We found that the equilibrium level of selfing was similar if F-loci are present (*p_f_* = 0.25) or absent (*p_f_* = 0), even though deleterious fixation rates are higher if F-loci are present.

Highly selfing species appear to be prone to extinction though we lack a detailed understanding of how this occurs. Our work indicates fluctuations in the strength of purifying selection can be an important contributor to this process. The results presented may underestimate the relative importance given that we observe stronger effects under lower mutation rates (Fig. 1, S13, S14). Even if rare in the genome, loci experiencing fluctuating selection make a disproportionately larger contribution to mutation accumulation and fitness decline than loci under constant selection. Moreover, mutation accumulation at such loci may make selfing species particularly susceptible to extinction to rare environmental conditions where the functions of such loci are particularly important. Although detecting and measuring the fitness impact of loci under fluctuating selection will be empirically challenging, our results demonstrate a plausible parameter space where fluctuating selection can strongly impact the mutation accumulation in highly selfing species.

